# Healthy cells functionally present TAP-independent SSR1 peptides: implications for selection of clinically relevant antigens

**DOI:** 10.1101/2020.06.11.146449

**Authors:** Antonius A. de Waard, Tamara Verkerk, Kelly Hoefakker, Dirk M. van der Steen, Marlieke L.M. Jongsma, Sophie Bliss, Arnoud H. de Ru, Peter A. van Veelen, Marieke Griffioen, Mirjam H.M. Heemskerk, Robbert M. Spaapen

## Abstract

Tumors with an impaired transporter associated with antigen processing (TAP) present several ER-derived self-antigens on HLA class I (HLA-I) which are absent on healthy cells. Selection of such TAP-independent antigens for T cell-based immunotherapy should include analysis of their expression on healthy cells to prevent therapy-induced adverse toxicities. However, it is unknown how the absence of clinically relevant antigens on healthy cells needs to be validated. Here we monitored TAP-independent antigen presentation on various healthy cells using a new toolbox consisting of a T cell clone recognizing a TAP-independent SSR1-derived antigen. We found that most but not all healthy cells present this antigen under normal and inflammatory conditions, indicating that TAP-independent antigen presentation is a variable phenomenon. Our data emphasize the necessity of extensive testing of a wide variety of healthy cell types to define clinically relevant TAP-independent antigens that can be safely targeted by immunotherapy.

## Introduction

HLA class I (HLA-I) (neo)antigen presentation by tumor cells can induce activation of CD8^+^ cytotoxic T cells (Neefjes et al., 2011). This natural immunological pressure forces various tumors to downmodulate HLA-I antigen presentation through genomic or epigenetic alterations (Campoli and Ferrone, 2008; Khong and Restifo, 2002). In addition, T cell based immunotherapies increase mutation rates in genes associated with antigen presentation such as *B2M, tapasin, TAP1* and *TAP2* (Gettinger et al., 2017; Jiang et al., 2010; Restifo et al., 1996; Sade-Feldman et al., 2017).

HLA-I is generated and folded in the ER, assisted by several lectin chaperones and stabilized by the beta-2 microglobulin (B2M) light chain, enabling the loading of small peptides through the collaborative effort of tapasin, calreticulin and ERp57 (Rock et al., 2016). Conventional peptide generation is mediated by the proteasome in the cytosol. These peptides are transported into the ER by the heterodimeric transporter associated with antigen processing (TAP) complex.

Alterations in the antigen presentation machinery affect the composition of the peptide repertoire presented by HLA-I (van Hall et al., 2006). Perhaps the largest impact on the repertoire is the functional disruption of the TAP transporter (hereafter referred to as TAP-deficient), interrupting the supply of cytosolic peptides. Without TAP, HLA-I presented peptides are either derived from ER-resident proteins or enter the ER through routes other than the canonical TAP pathway (Oliveira and van Hall, 2015). Several of these so-called TAP-independent peptides are not presented on healthy TAP-proficient cells, therefore representing a potential specific immunotherapeutic target on TAP-deficient tumors (van Hall et al., 2006). Since studies targeting TAP-independent peptides in mice show low toxicity, the first proof-of-concept study targeting TAP-independent peptides in patients with non-small cell lung cancer will soon be initiated (Brolsma, 2019; Doorduijn et al., 2018). Potential target antigens for such therapy were recently identified in humans by comparing the HLA-I presented peptidome eluted from TAP-deficient and -proficient cells (Marijt et al., 2018). Because a considerable proportion of TAP-independent peptides can also be presented by at least some TAP-proficient cells (Man et al., 1992; Weinzierl et al., 2008), clinical targeting of TAP-independent antigens potentially involves significant safety and efficacy risks. Although caution is needed in designating peptides as solely being presented on TAP-deficient cells, efforts are largely lacking to validate that antigens are not at all presented by any healthy cell type with native TAP expression.

To define a validation roadmap, it would be instrumental to have a tool - such as a defined T cell clone - to monitor functional TAP-independent peptide presentation both on tumor and healthy cells. However, isolation of T cells targeting TAP-independent antigens on healthy cells may be challenging since they are likely deleted during thymic development (Takaba and Takayanagi, 2017). Therefore, human T cells that recognize a molecularly defined TAP-independent antigen on healthy cells have not been identified to date.

To bypass potential thymic deletion issues, we here investigated TAP-independent antigen recognition on healthy cells using CD8^+^ T cell clones derived from allogeneic repertoires (Amir et al., 2011; Van Bergen et al., 2010). We identified that the cognate antigen of one of these clones, derived from the ER-resident protein SSR1, is presented both by TAP knockout (KO) and TAP-proficient tumor cells. This ubiquitously expressed antigen is effectively presented under normal and inflammatory conditions on several but not all healthy primary cells, indicating that TAP-independent antigen presentation is a variable phenomenon. Thus, a broad healthy cell expression analysis of TAP-independent targets improves the safety profile of their immunotherapeutic application.

## Results

### The protein SSR1 encodes a TAP-independent peptide

To study the effect of the TAP transporter on the functional presentation of antigens, we genetically knocked out TAP1 in human HAP1 cells. After lentiviral introduction of CRISPR/Cas9 machinery targeting the first exon of TAP1, 20% of the transduced cells still displayed surface HLA-I levels comparable to wild type cells (Figure 1A and S1). In order to prevent interference of remaining wild type cells in T cell coculture assays, we generated five clonal TAP1 KO cell lines which showed decreased HLA-I surface expression (Figure 1B). In addition, western blot analysis confirmed a complete lack of TAP1 expression in clone #1 (de Waard et al., 2020). Sequence analyses showed a lack of at least one amino acid in the first transmembrane domain of TAP1 in each clone (Figure S2A and B) (Oldham et al., 2016).

**Figure 1.**
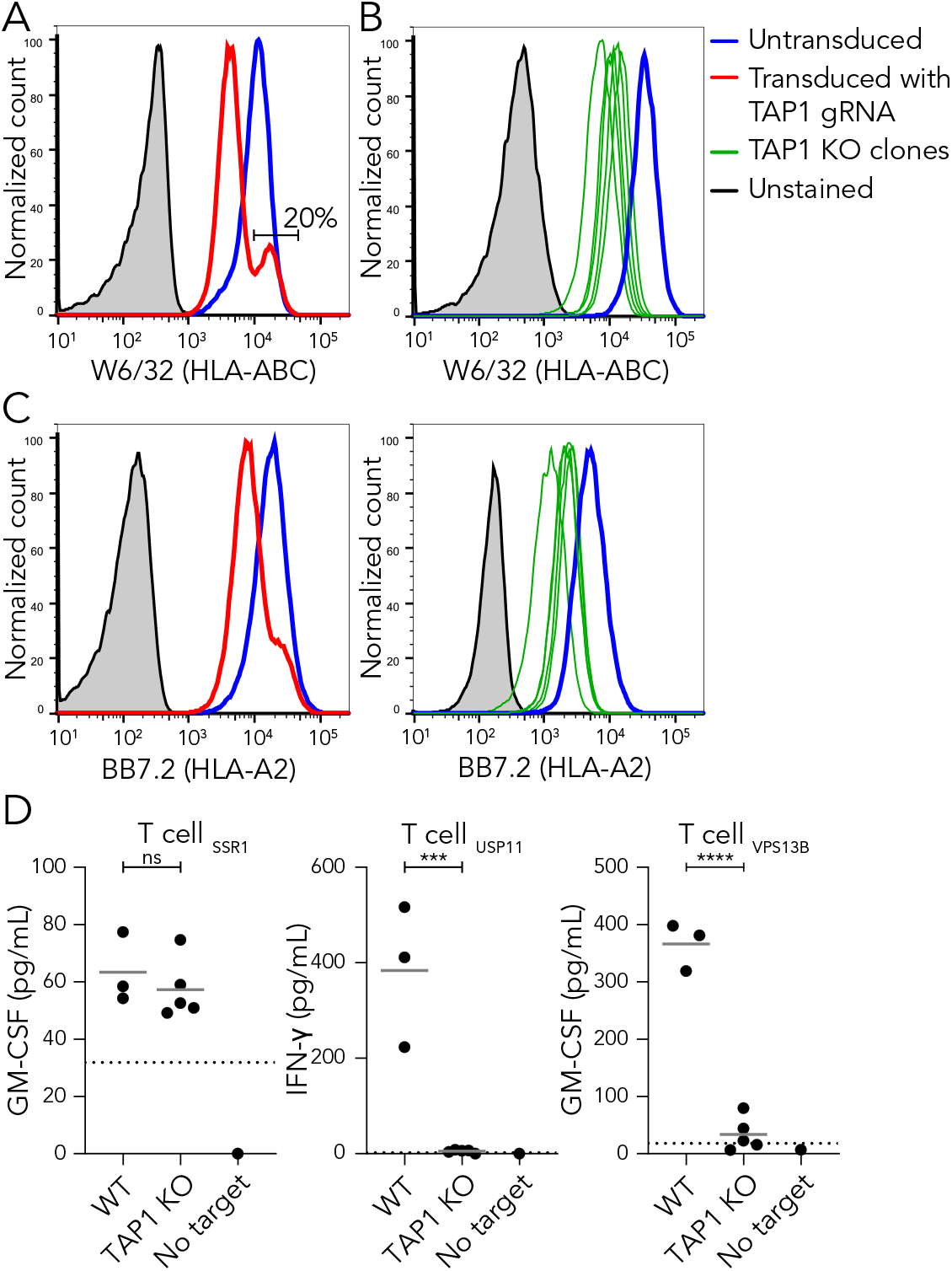
The protein SSR1 encodes a TAP-independent peptide. (**A**) HAP1 cells were transduced with a TAP1-specific gRNA and analyzed by flow cytometry using the pan-HLA antibody W6/32. Untransduced (blue), transduced (red) and unstained cells (grey). See Figure S1 for gating strategy. (**B**) Five monoclonal cell lines were generated from the polyclonal TAP1 KO cell line in (A) and analyzed by flow cytometry. Wild type (blue), TAP1 KO clones (green lines), unstained cells (grey). See Figure S2 for sequencing details. (**C**) Same as (A) and (B) but using the HLA-A2-specific antibody BB7.2. (**D**) HLA-A2-restricted SSR1-, USP11- or VPS13B-specific T cells were cocultured with three separate batches of HAP1 wild type or each of the five clonal TAP1 KO cell lines (B). Culture supernatant was used for indicated cytokine ELISAs. Each datapoint represents the average of triplicate cultures. Dotted lines represent the lower detection limit of the ELISA. Student’s t-test, ***p<0.001, ****p<0.0001, ns is not significant.

To analyze functional presentation of antigens in the absence of the TAP transporter, we utilized several alloreactive CD8^+^ T cell clones that were previously isolated from patients following allogeneic stem cell transplantation (Amir et al., 2011; Van Bergen et al., 2010). These T cell clones each recognize a defined antigen derived from the cytosolic proteins USP11 or VPS13B, or from the ER-localized transmembrane protein SSR1 (Koike and Jahn, 2019; Pfeffer et al., 2017; Zhou et al., 2017). Since these antigens are presented by HLA-A2, we first confirmed that this specific allele was affected in the TAP1 KO cells (Figure 1C). Subsequent coculture of T cells with HAP1 target cells showed that USP11- and VPS13B-specific T cells recognized wild type, but not TAP-deficient cells (Figure 1D, right panels). Conversely, SSR1-specific T cells were activated by the TAP-deficient cells demonstrating that the SSR1 peptide is processed independently of TAP (Figure 1D, left panel). Strikingly, HAP1 wild type cells were capable of activating SSR1-specific T cells to a similar level as TAP-deficient cells, showing that the SSR1 peptide can be properly presented at the cell surface even in the presence of the TAP transporter (Figure 1D). Thus, the SSR1 peptide represents a new model antigen to further evaluate TAP-independent antigen presentation by healthy cells.

### The SSR1 signal peptide contains the TAP-independent SSR1 antigen

The epitope recognized by the SSR1-specific T cell clone is polymorphic in the population as dictated by a nonsynonymous SNP (rs10004) encoding for either a serine or leucine (SSR1-S of SSR1-L) (Van Bergen et al., 2010). A 14-mer peptide containing this polymorphism (VLFRGGPRG<u>S</u>LAVA) was previously shown to be presented by HLA-A2 and recognized by SSR1-specific T cells, albeit with low avidity (Bijen et al., 2018). As more specific details on functionally presented SSR1 antigens may aid in defining TAP-independent peptide processing process, we next investigated the HLA-I mediated presentation and recognition of additional peptides covering the SNP. *In silico* HLA-A2-binding predictions of SSR1 peptides covering the SNP failed to designate strong binders (Table S1). Since predictions for this SSR1 epitope can be inaccurate (Bijen et al., 2018), we biochemically determined whether SSR1 peptides other than the 14-mer are presented by HLA-A2 positive cells. Using LC-MS/MS we analyzed eluted peptides from HAP1 wild type cells (de Waard et al., 2020), and from two HLA-A2 positive primary AML samples that genetically encode the SSR1-S peptide. In first instance, we automatically retrieved only SSR1-L peptides (13-mer and 14-mer) from the AML samples (Table 1). In addition, using the MS2 spectrum of a synthetic SSR1-S 14-mer peptide as reference, we identified this peptide in all three samples (Table 1).

**Table 1.** Immunopeptidome analyses of cells genetically positive for SSR1-S. SSR1-S and SSR1-L candidate peptides were eluted from HAP1 cells and two primary AMLs. Affinities for HLA-A*02:01 were predicted using NetMHC4.0.

| Candidate peptide | Identified on | Genotype rs10004 | Predicted affinity for HLA-A*02:01 (nM) |
| --- | --- | --- | --- |
| VLFRGGPRG**S**LAVA | HAP1,<br>AGP2375,<br>SWD2031 | S/S,<br>S/L,<br>S/L | 1403 |
| VLFRGGPRG**L**LAVA | AGP2375 | S/L | 649 |
| VLFRGGPRG**L**LAV | SWD2031 | S/L | 236 |

To obtain more data on presented SSR1-derived peptides, we analyzed the peptidome from additional HLA-A2 positive primary tumor samples (n=9) and cell lines (n=6). As both SSR1-S and SSR1-L are presented at similar levels by HLA-A2 positive cells (Bijen et al., 2018), we further focused on identifying length variants of the SSR1-L peptide. The analysis yielded four length variants of the same core peptide (<u>VLFRGGPRGLL</u>*AVA*) covering the polymorphic residue (Table 2). The fact that candidate peptides were derived from one TAP-deficient (T2) cell line and many presumably TAP-proficient cells supported our findings that presentation of the SSR1 antigen is independent of TAP, but not restricted to cells lacking TAP function. Coculture of SSR1-specific T cells with S-antigen-negative T2 cells loaded with decreasing amounts of synthetic candidate peptides showed superior responses against the 14-mer peptide (Figure 2A). These data establish the 14-mer peptide as the cognate epitope of the SSR1-specific T cell clone.

**Figure 2.**
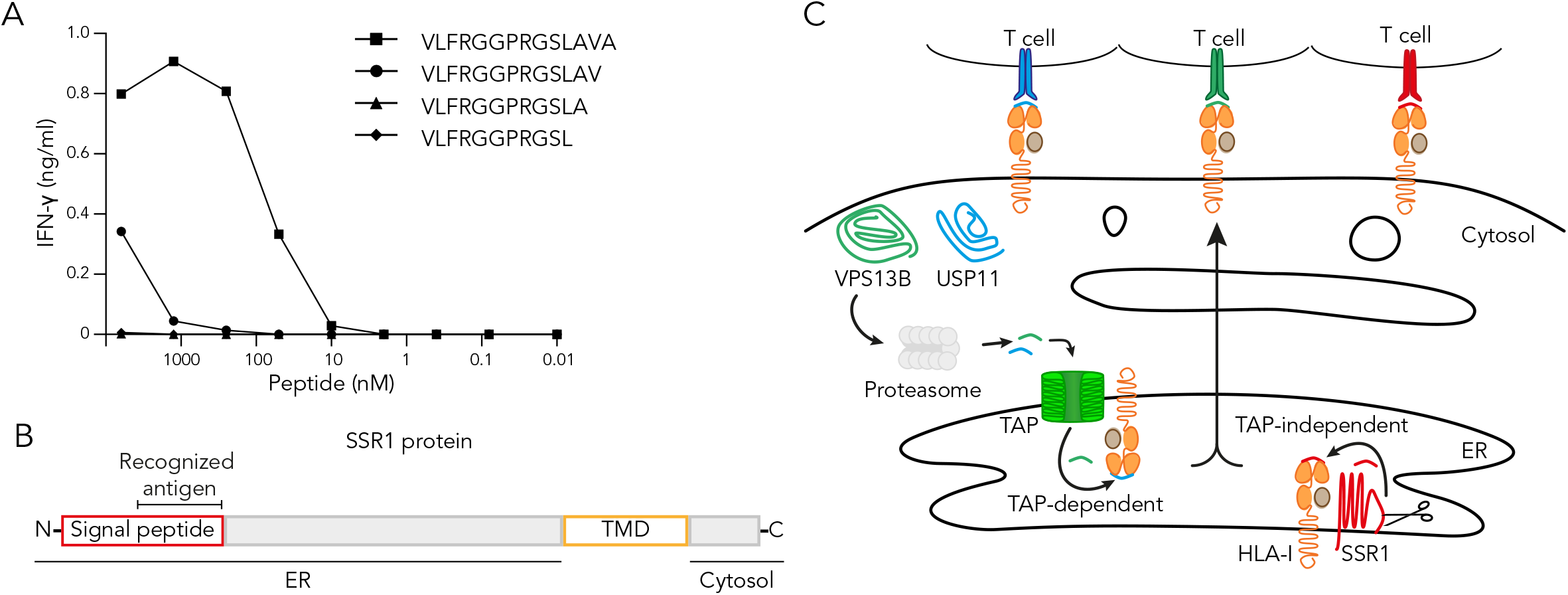
The SSR1 signal peptide contains the TAP-independent SSR1 antigen. (**A**) T2 cells, loaded with different amounts of synthetic length variants of the SSR1-S peptide (peptide sequences in Table 2), were cocultured with SSR1-specific T cells after which culture supernatant was used for IFN-γ ELISA. Each datapoint represents a single measurement. (**B**) Summary of the location of the antigenic peptide in the SSR1 protein. TMD, transmembrane domain (yellow). (**C**) Graphic summary of subcellular localization of the three parental proteins studied here and the presumed pathway of their antigenic peptides for presentation on HLA-I to their cognate T cells.

**Table 2.** Immunopeptidome analysis reveals seven candidate peptides covering the SSR1 SNP (rs10004) SSR1-L candidate peptides eluted from different primary tumors (anonymized patient numbers indicated) and cell lines. Affinities for HLA-A*02:01 were predicted using NetMHC4.0 for both SSR1-L and SSR1-S of rs10004.

| Candidate peptide | Identified from |  |  | Predicted affinity for HLA-A*02:01 (nM) per variant |  |
| --- | --- | --- | --- | --- | --- |
|  | Primary tumor | Cell line | TAP-deficient cell line | L | S |
| VLFRGGPRG**LL** | - | - | T2 | 4673 | 5524 |
| VLFRGGPRG**LLA** | OvaL10 | UM9+ A2 | - | 5795 | 10082 |
| VLFRGGPRG**LLAV** | SWD2031 | - | - | 236 | 628 |
| VLFRGGPRG**LLAVA** | AGP2375,<br>CDT4512,<br>KYE9115,<br>OvaL10,<br>OvaL23,<br>GBZ10197,<br>GPQ3361,<br>VDL6711,<br>ALY1143 | U266,<br>TMD8+A2,<br>COV413b,<br>U2G10196,<br>UM9+A2 | T2 | 649 | 1403 |

The location of an antigen within a transmembrane protein determines the molecular path of presentation, including the requirement for TAP transportation after proteolytic degradation (Oliveira and van Hall, 2013). SSR1 is a type-I transmembrane protein with an intra-ER domain of 189 amino acids and a cytosolic domain of 58 amino acids (Pfeffer et al., 2017). To gain insight into the presentation route of the SSR1 antigen, we analyzed the subcellular localization of the recognized peptide. The 14-mer peptide is located near the N-terminus of the SSR1 protein, spanning part of the H- and the complete C-region of the signal peptide as predicted by Phobius (Kall et al., 2007), and is likely to be C-terminally liberated in the ER by the signal peptidase complex (Figure 2B) (Voldby Larsen et al., 2006; Weinzierl et al., 2008). In summary, the peptides derived from USP11 and VPS13B are processed in the cytosol and require TAP to enter the ER and bind to HLA-I, whereas the SSR1 peptide is readily localized in the ER and likely processed by local proteases before being loaded on HLA-I (Figure 2C).

### TAP-independent peptide presentation is functional in healthy donor cells

To investigate the capacity of non-transformed cells to present TAP-independent peptides, we determined the reactivity of the SSR1-specific T cell clone against healthy donor material. Monocytes, T cells and B cells, the main constituents of peripheral blood mononuclear cells (PBMCs), express SSR1 according to publicly available RNA-seq data (Figure 3A). Therefore, we isolated PBMCs from three HLA-A2 positive healthy donors and genotyped rs10004 to evaluate whether these donors were carrying the antigen recognized by the SSR1-specific T cell clone. Two donors (A and B) were heterozygous for rs10004 and thus positive for the SSR1 epitope, and one donor (C) was homozygous negative (Figure 3B). Coculture of our T cell clones with fresh PBMCs from the three donors failed to elicit a detectable response of SSR1-specific T cells, while the positive control USP11- and VPS13B-specific clones were activated (Figure 3C). In contrast, monocytes isolated from the genetically positive PBMCs induced an SSR1-directed T cell response (Figure 3D). We then interrogated T cell reactivity against human umbilical cord endothelial cells (HUVECs), as a non-hematopoietic healthy cell type that expresses SSR1 (Figure 3E). SSR1-specific T cells responded to selected HLA-A2 positive, SSR1-antigen positive HUVECs, but not to HLA-A2 negative control HUVECs (Figure 3F and G). Together, these data demonstrate that presentation of the TAP-independent SSR1-antigen is functional in multiple, but not all healthy TAP-proficient cells.

**Figure 3.**
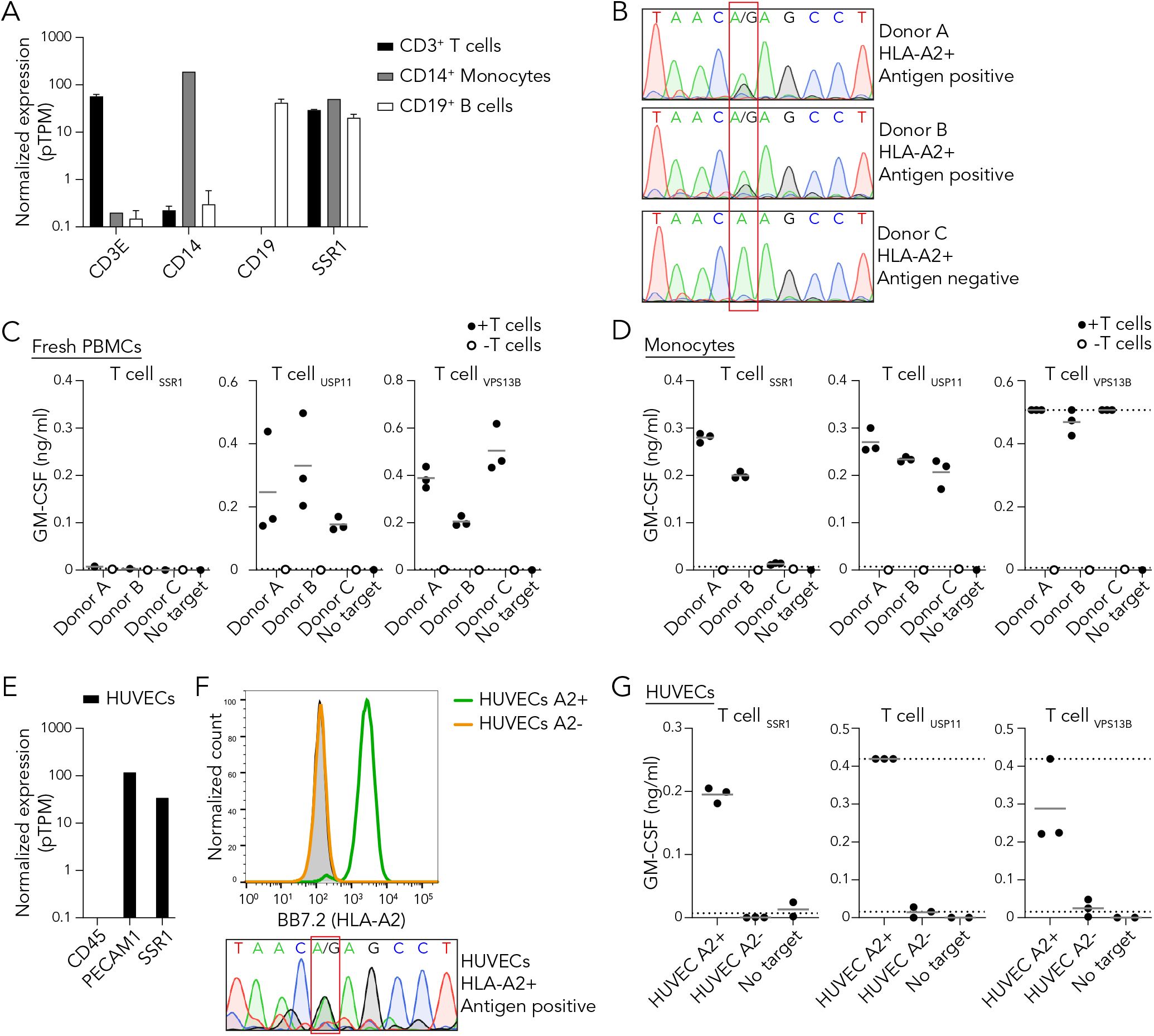
TAP-independent peptide presentation is functional in healthy donor cells. (**A**) Normalized expression of SSR1 and control CD14, CD19 and CD3E per depicted PBMC subset using the RNA HPA blood cell gene data. (**B**) Sanger sequencing of the SSR1 SNP rs10004 of three HLA-A2 positive healthy donors, the position of the SNP is highlighted by the red box and consequence for antigen status is depicted. (**C**) T cell coculture with PBMCs from three healthy donors (B), culture supernatant was analyzed by GM-CSF ELISA. Results from triplicate cultures are shown. Dotted lines represent the detection limits of the ELISA. (**D**) As in (C) but using monocytes as target cells. (**E**) Expression analysis of SSR1 and control CD45 and PECAM1 in TERT2 immortalized HUVECs using the RNA HPA cell line gene data. (**F**) Flow cytometric analysis of HLA-A2 expression by HUVECs. HLA-A2 positive FACS sorted (green), HLA-A2 negative FACS sorted (orange) populations and unstained HUVECs (grey). Lower panel depicts rs10004 sequence of the HLA-A2 positive sorted HUVECs. (**G**) T cell coculture with HLA-A2 positive and negative HUVECs (F). Culture supernatant was analyzed by GM-CSF ELISA. Each dot represents an individual measurement of triplicate cultures.

### TAP-independent peptides are presented under inflammatory conditions

Patients receiving immunotherapy regularly experience severe inflammatory reactions and cytokine storm, which may also be the case after immunotherapeutic targeting of TAP-independent peptides (Gangadhar and Vonderheide, 2014; Hay et al., 2017; Henden and Hill, 2015). Inflammatory cytokines such as IFN-γ enhance expression of various components of the HLA-I pathway including the TAP transporter, but their influence on the expression and presentation of the antigens studied here is unknown (Jongsma et al., 2019; Ma et al., 1997). Therefore, we first analyzed the change in expression of SSR1, USP11 and VPS13B after 24 hours of IFN-γ treatment using microarray datasets from the Interferome database (Rusinova et al., 2012). In comparison to the reference gene *PPIA* (Riemer et al., 2012), the expression of *SSR1, USP11* and *VPS13B* were unaffected by IFN-γ treatment whereas *TAP1* and *TAP2* expression were increased (Figure 4A). Additionally, proteome analysis after 48 hour IFN-γ stimulation revealed no significant change in SSR1 and USP11 protein levels while TAP1 and TAP2 increased eight-fold (Figure 4B) (Megger et al., 2017).

**Figure 4.**
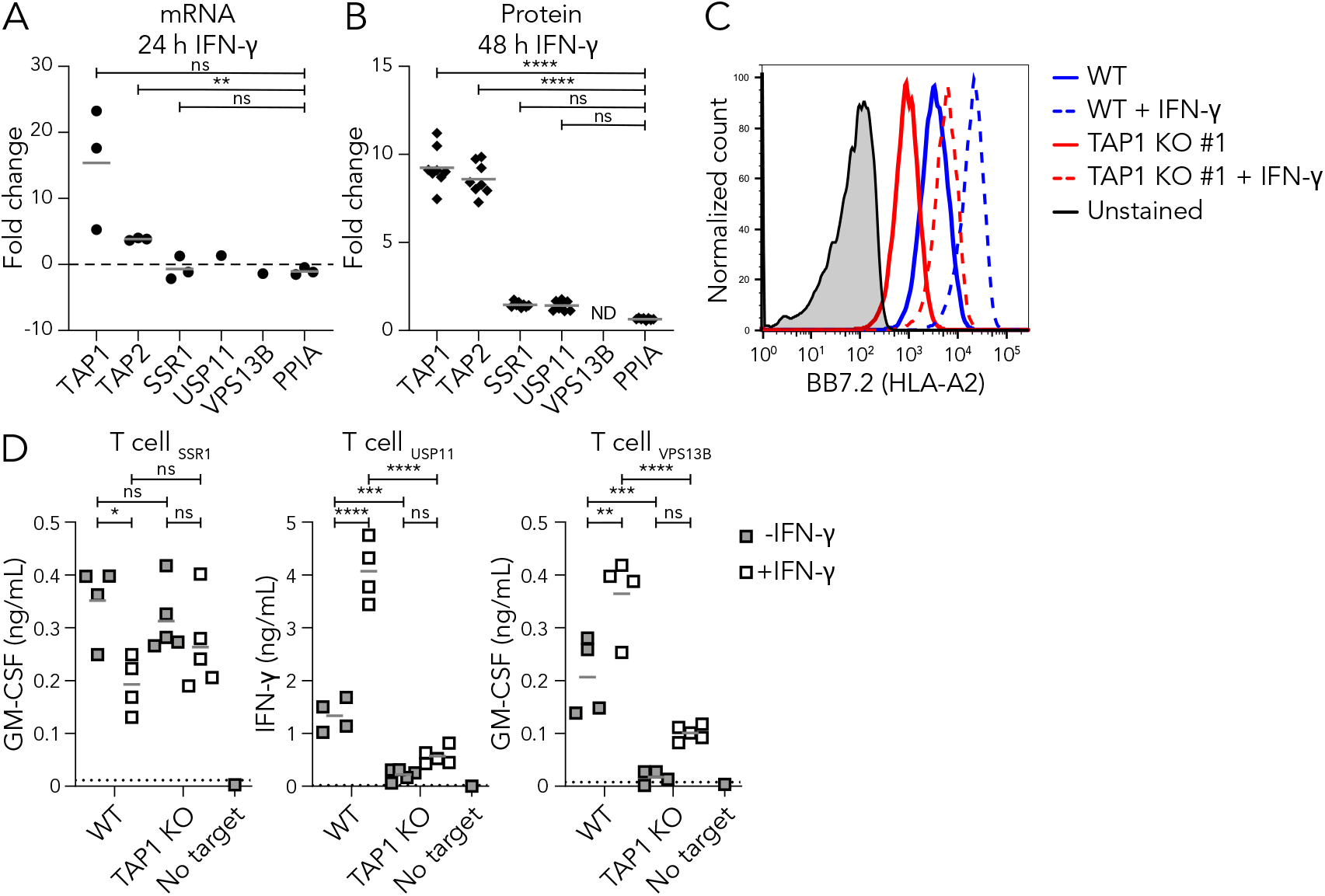
TAP-independent peptides are presented under inflammatory conditions. (**A**) *TAP1, TAP2, SSR1, USP11* and *VPS13B* mRNA expression after 24 hours of IFN-γ stimulation was analyzed using three publicly available datasets and compared to the reference gene *PPIA*. (**B**) TAP1, TAP2, SSR1, USP11 and VPS13B protein expression after 48 hours of IFN-γ stimulation was analyzed using publicly available MS data and compared to the reference protein PPIA (Megger et al., 2017). ND is not detected. (**C**) HAP1 wild type and TAP1 KO cells were treated with IFN-γ for 48 hours and surface HLA-A2 upregulation was measured by flow cytometry. Unstimulated (solid) or stimulated (dashed) wild type (blue) and TAP1 KO #1 (red) cells are shown, including unstimulated unstained wild type cells (grey). (**D**) T cell coculture with unstimulated or IFN-γ stimulated wild type or TAP1 KO clones (same as in Figure 1B). Culture supernatants were analyzed for the presence of cytokines using cytokine ELISA. Dotted lines represent the lower and upper detection limits of the ELISA. Repeated measures one-way ANOVA followed by Dunnett’s (A), ordinary one-way ANOVA followed by Dunnett’s (B) or by Sidak’s (D) multiple comparisons test. *p<0.05, **p<0.01, ***p<0.001, ****p<0.0001, ns is not significant.

To examine the effects of inflammatory conditions on TAP-independent peptide presentation, we stimulated HAP1 wild type and TAP1 KO cells for 48 hours with IFN-γ, which resulted in increased HLA-A2 surface expression on both cell types (Figure 4C). These stimulated cells and their unstimulated controls were then exposed to the different T cell clones. The IFN-γ-induced upregulation of classical antigen presentation on wild type cells was characterized by an increased T cell response against control USP11 and VPS13B peptides (Figure 4D, right panels). These IFN-γ stimulated wild type cells also presented significant amounts of SSR1 peptide (Figure 4D, left panel). In addition, the SSR1 peptide presentation in the absence of the TAP transporter was similar after IFN-γ stimulation (Figure 4D). These results indicate that this TAP-independent antigen presentation pathway is largely unaffected by an inflammatory IFN-γ stimulus on an immortal model cell line.

### Activated healthy cells functionally present TAP-independent peptides

To evaluate whether healthy tissues present TAP-independent peptides under inflammatory conditions similar as induced by immunotherapy, we evaluated recognition of the TAP-independent SSR1 antigen on activated healthy cells. Although functional SSR1 antigen presentation by unstimulated PBMCs was negligible (Figure 3C), we observed a significant SSR1-specific T cell response against IFN-γ treated PBMCs from the SSR1 antigen positive donors A and B, but not against the antigen-negative donor C (Figure 5A). In order to test the TAP-independent antigen presentation in a broader inflammatory context, we activated B cells of the same three donors using CpG and a BCR-stimulus (Figure 5B). These activated B cells were efficiently recognized by SSR1-specific T cells, further substantiating that an inflammatory environment allows for TAP-independent antigen presentation in healthy cells. Finally, the SSR1-specific T cells were also reactive to non-immune cells (HUVECs) pre-exposed to IFN-γ (Figure 5C). Together, our data demonstrate that healthy TAP-proficient cells can efficiently present TAP-independent peptides under steady state and inflammatory conditions.

**Figure 5.**
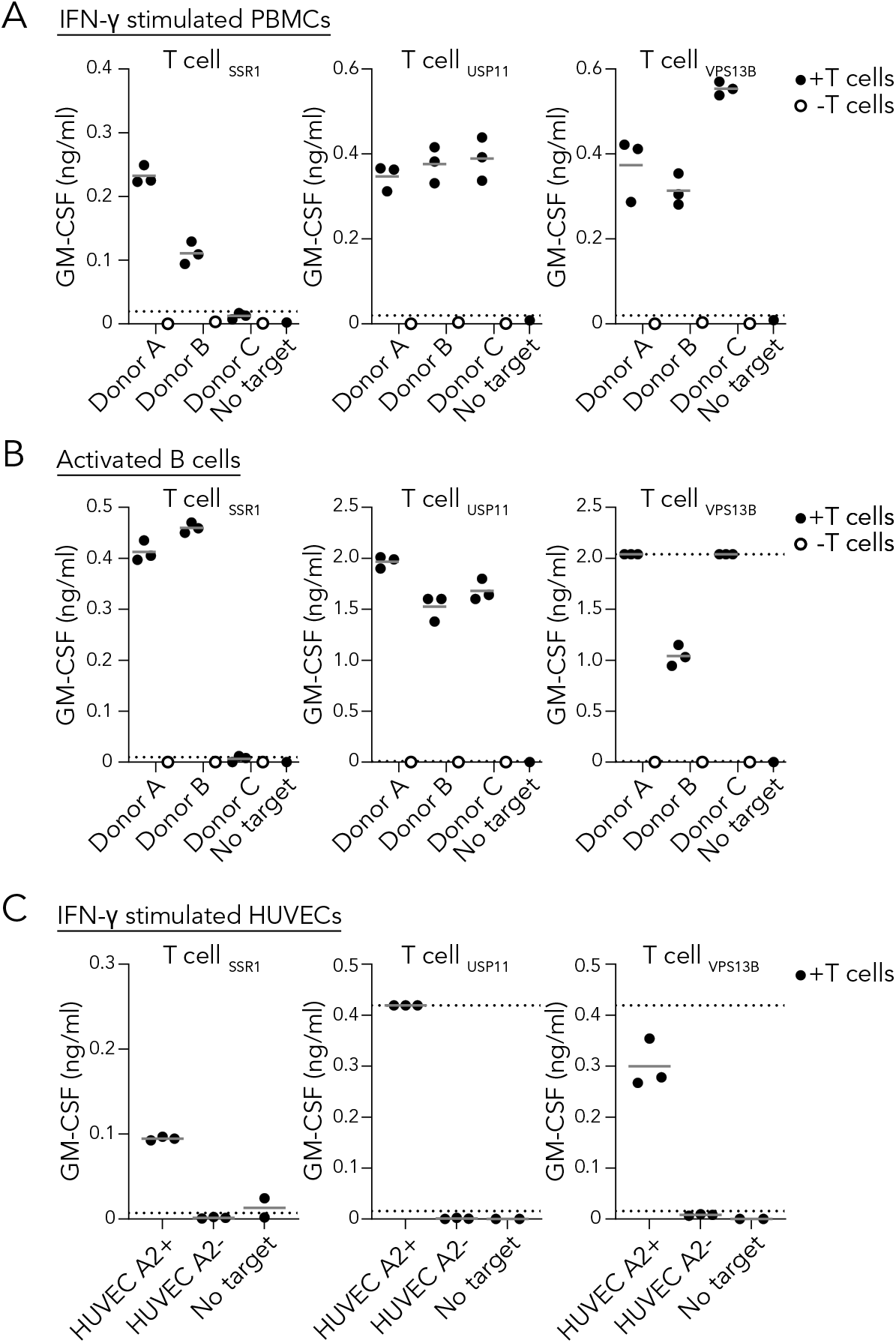
Activated healthy cells functionally present TAP-independent peptides. (**A**) T cell coculture with healthy donor PBMCs activated for 3 days with IFN-γ. (**B**) T cell coculture with healthy donor B cells activated using CpG and B cell receptor crosslinking antibodies. (**C**) T cell coculture with HLA-A2 positive and negative HUVECS activated with IFN-γ. Culture supernatants were analyzed using GM-CSF ELISA. Results from triplicate cultures are shown. Dotted lines represent lower and upper detection limit of the ELISAs.

## Discussion

Tumor specific abrogation of the antigen presentation machinery can provide opportunities for new therapeutic approaches, such as targeting (neo)antigens that are presented only in the case of a defective TAP transporter (Marijt and van Hall, 2020; Oliveira and van Hall, 2013; van Hall et al., 2006). In our current study, we identified that an immunogenic T cell epitope originating from the ER-resident protein SSR1 is processed independently of TAP. Using an SSR1-specific T cell clone, we provide evidence that TAP-independent presentation is variable on various healthy cell types under resting and inflammatory conditions. Thus, to ensure the safety and efficacy of a therapy against TAP-independent antigens, our data imply that it is vital for every individual target to map the degree of presentation on healthy tissue.

Therapeutic targeting of non-mutated TAP-independent antigens has potential for broad clinical application, since these antigens are shared among patients with TAP-deficient tumors (Oliveira et al., 2013; van Hall et al., 2006; Voldby Larsen et al., 2006). Such immunotherapies will utilize either vaccination strategies or adoptive transfer of specific T cells. A strict prerequisite for a vaccination strategy to be effective is the lack of expression of the targeted antigen(s) on healthy cells, because otherwise specific T cells will be functionally impaired or deleted during thymic development (Doorduijn et al., 2016; Takaba and Takayanagi, 2017). Importantly, such immune tolerance inherently provides a safety margin for clinical studies in which presentation of targeted antigens would unexpectedly be not restricted to TAP-deficient cells. To increase the chance on raising an effective immune response against TAP-independent antigens, vaccination with TAP-silenced DCs could be a potential strategy (Marijt et al., 2019). Of note, there are multiple reports that in humans vaccination strategies against antigens presented on healthy tissue can break tolerance and induce severe autoimmunity, so caution is warranted (Ludewig et al., 2000; Sultan et al., 2017). The alternative for immunotherapeutic vaccination is adoptive T cell transfer, which can induce a fast anti-tumor response. A combination of adoptive transfer of T cells specific for a tumor-restricted TAP-independent antigen with repeated long peptide vaccination proved effective against murine TAP-deficient tumors (Doorduijn et al., 2016). However, expression of a target antigen by only a subset of healthy cells can induce lethal autoimmune toxicity (Cameron et al., 2013; Morgan et al., 2013; Yee et al., 2000). Our study emphasizes that TAP-independent peptides may be presented by a subset of TAP-proficient cells. Thus, before clinical targeting of TAP-independent antigens, it is crucial to apply a thorough selection with the goal to exclude antigens that are presented on healthy cells.

To optimally select target antigens, mechanistic insights into the intracellular process of peptide selection are required. Current knowledge on how certain TAP-independent peptides fail to be selected for presentation in the presence of TAP is limited. Hypotheses are based on the peptide affinity for HLA-I, its abundance in the ER, and the distance between generation site and peptide-receptive HLA-I molecules (Durgeau et al., 2011; Oliveira et al., 2011; Oliveira and van Hall, 2013). Our human TAP-independent SSR1 antigen and its cognate T cells represent a unique toolset to further investigate existing and novel hypotheses. In support of the affinity hypothesis, the 14-mer SSR1 peptide, which has an HLA-A2 affinity well below the generally accepted threshold for presentation of 500 nM (78 nM), is selected for presentation in the presence of TAP (Bijen et al., 2018; Paul et al., 2013). However, several TAP-independent peptides that are not presented by a TAP-proficient model cell line have a strong predicted HLA-I binding, suggesting that peptide presentation decisions are not solely dictated by HLA-I affinity (Marijt et al., 2018). The second hypothesis suggests that highly abundant peptides pumped into the ER by TAP may be presented at the expense of less abundant TAP-independent peptides (Oliveira et al., 2011). In line with this hypothesis, boosting TAP-mediated competitor peptide supply into the ER by IFN-γ seemed to decrease TAP-independent SSR1 antigen presentation on wild type cells. In contrast, a reduction in competitor peptide abundance by depleting TAP from wildtype cells failed to enhance SSR1 peptide presentation, suggesting that also other factors play a role. Because the general abundance of TAP-dependent peptides positively correlates with the expression level of TAP (Boegel et al., 2018; Durgeau et al., 2011), the presentation of TAP-independent peptides may be more efficient on cells with low TAP expression, as long as sufficient peptide-receptive HLA-I heavy chains are available. Thus, presentation of a putative TAP-independent target antigen depends at least on the expression of its encoding gene, TAP, HLA-I, and also on its affinity and the amount of competitor peptides. Therefore, safe TAP-independent clinical target definition requires assessment of a broad panel of resting and inflamed healthy cells, including cells with differential TAP and antigen-encoding gene expression. As shown here, PBMC-derived cells and HUVECS can be included in such a panel, but also other healthy human cells such as but not limited to fibroblasts, keratinocytes, melanocytes, differentiated induced pluripotent stem cells (iPSCs) or organoids from different organs.

Taken together, the TAP-independent, polymorphic SSR1 antigen provides novel opportunities to improve our understanding of TAP-independent peptide presentation through T cell- and MS-based experiments on healthy and TAP-deficient cells. Our data warrant testing presentation of candidate TAP-independent target antigens on a broad panel of healthy cells before clinical application.

## Limitations of the study

Although we provide a direct proof of principle of TAP-independent antigen presentation by TAP-proficient cells, we have only tested a limited number of healthy donor-derived cells. Panels focusing on validating TAP-independent antigens for immunotherapy should rely on a broader panel of cell types derived from other vital tissues and organs. A technical limitation of this study is that none of the obtained TAP1 KO clones had a frameshifting mutation on both chromosomes. Although we showed absence of TAP1 by immunoblot (de Waard et al., 2020), a decrease in surface HLA-I, and a lack of presentation of cytosolic antigens, we cannot rule out residual TAP function in our model cells. The facts that the peptide is derived from a signal sequence in the ER, and that we and others eluted the SSR1 peptide from TAP-deficient T2 and 721.174 cell lines respectively (Weinzierl et al., 2008), provide further support of the TAP-independence of the SSR1 antigen.

## Acknowledgments

The authors thank the Sanquin Research Facility for their assistance with flow cytometry and Janine Arts and Dr. Jaap van Buul for providing HUVECs and assistance with their culture. This work was supported by the Netherlands organization for scientific research (NWO-VENI 016.131.047; R.S.), KWF Alpe d’HuZes (Bas Mulder Award 2015-7982; R.S.), the Landsteiner Foundation for Blood Transfusion Research (LSBR fellowship 1842F; R.S.) and an investment Grant NWO Medium (91116004, partly financed by ZonMw) to P.A.v.V.

## Author Contributions

Conceptualization and design: A.W. and R.S.; Data acquisition, analysis and interpretation: A.W., T.V., K.H., D.S., M.J., S.B., A.R., R.S.; Resources and discussion: M.G., M.H., P.V., R.S.; Supervision and conceptual discussion: R.S.; Writing: A.W., R.S.; Editing: A.W., T.V. and R.S..

---

**Figure S1.**
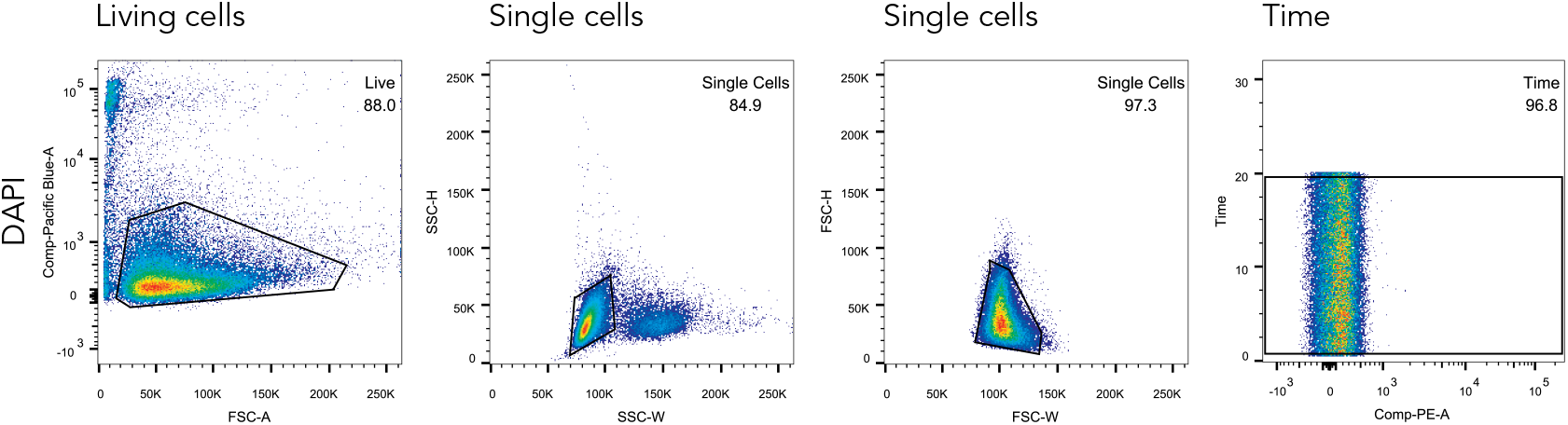
Gating strategy for flow cytometry. For all flow cytometric analyses, live cells were gated on DAPI negativity followed by two gates for single cells using SSC-H vs SSC-W and FSC-H vs FSC-W. Lastly, any potential artifacts induced by clogs were excluded from analysis by gating on time vs PE.

**Figure S2.**
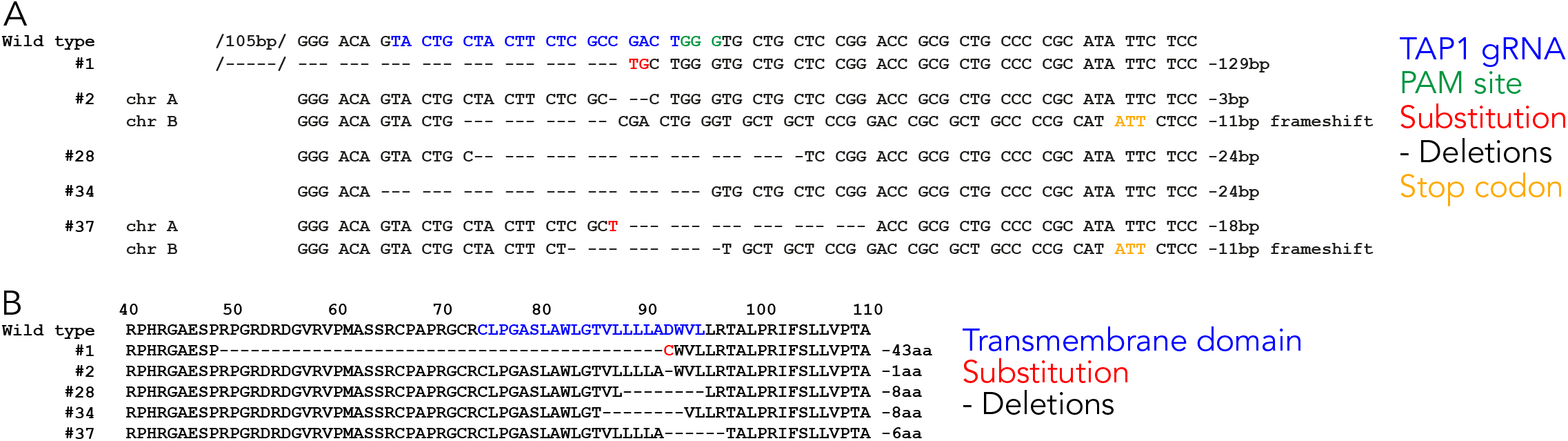
HAP1 TAP1 KO clone characteristics (Related to Figure 1) (**A**) Summary of TAP1 gene editing in each TAP1 KO clone. Indicated are the gRNA targeting sequence (blue) with PAM site (green), insertions (red), deletions (dashes) and premature stop codons (orange). Clone #1 was previously described (de Waard et al., 2020). (**B**) Summary of the targeted part of the TAP1 amino acid sequence with the genome editing result depicted for each clone. Numbers above the sequence indicate the amino acid location in the wild type TAP1 protein. Transmembrane domain in wild type sequence (blue), amino acids lacking in TAP1 KO clones (dashes) and amino acid substitutions (red) are depicted.

**Table S1.** All theoretical 8-14mer peptides covering the SSR1 SNP. Affinities for HLA-A*02:01 of all theoretical SSR1-derived peptides covering the nonsynonymous SNP rs10004 predicted using NetMHC4.0.

|  | Predicted affinity for HLA-A*02:01<br>(nM) per variant |  |
| --- | --- | --- |
| **Peptide sequence** | **L** | **S** |
| FRGGPRGL | 36658 | 43601 |
| RGGPRGLL | 39127 | 41929 |
| GGPRGLLA | 38138 | 43187 |
| GPRGLLAV | 26994 | 33459 |
| PRGLLAVA | 35607 | 39522 |
| RGLLAVAQ | 34391 | 37964 |
| GLLAVAQD | 27129 | 40492 |
| LLAVAQDL | 6903 | 10021 |
| LFRGGPRGL | 30649 | 39466 |
| FRGGPRGLL | 31365 | 34114 |
| RGGPRGLLA | 29547 | 36909 |
| GGPRGLLAV | 16626 | 23894 |
| GPRGLLAVA | 26057 | 30604 |
| PRGLLAVAQ | 34960 | 38808 |
| RGLLAVAQD | 32556 | 35998 |
| GLLAVAQDL | 286 | 13769 |
| LLAVAQDLT | 5043 | 5748 |
| VLFRGGPRGL | 777 | 16308 |
| LFRGGPRGLL | 33415 | 35308 |
| FRGGPRGLLA | 26878 | 36619 |
| RGGPRGLLAV | 16823 | 23776 |
| GGPRGLLAVA | 26366 | 33402 |
| GPRGLLAVAQ | 32540 | 36539 |
| PRGLLAVAQD | 37729 | 40484 |
| RGLLAVAQDL | 1447 | 20810 |
| GLLAVAQDLT | 2684 | 18948 |
| LLAVAQDLTE | 21253 | 23246 |
| TVLFRGGPRGL | 7427 | 30851 |
| VLFRGGPRGLL | 4673 | 5524 |
| LFRGGPRGLLA | 32850 | 38768 |
| FRGGPRGLLAV | 18430 | 25942 |
| RGGPRGLLAVA | 29760 | 35464 |
| GGPRGLLAVAQ | 34513 | 38438 |
| GPRGLLAVAQD | 36423 | 39518 |
| PRGLLAVAQDL | 8766 | 30236 |
| RGLLAVAQDLT | 11515 | 29918 |
| GLLAVAQDLTE | 15158 | 31575 |

| Peptide sequence | L | S |
| --- | --- | --- |
| LLAVAQDLTED | 32609 | 34026 |
| ATVLFRGGPRGL | 8580 | 31521 |
| TVLFRGGPRGLL | 6482 | 7682 |
| VLFRGGPRGLLA | 5795 | 10082 |
| LFRGGPRGLLAV | 20279 | 27386 |
| FRGGPRGLLAVA | 23552 | 30074 |
| RGGPRGLLAVAQ | 32152 | 36590 |
| GGPRGLLAVAQD | 35448 | 39262 |
| GPRGLLAVAQDL | 7843 | 30371 |
| PRGLLAVAQDLT | 13387 | 30950 |
| RGLLAVAQDLTE | 15461 | 31263 |
| GLLAVAQDLTED | 16550 | 34127 |
| LLAVAQDLTEDE | 33202 | 34697 |
| PATVLFRGGPRGL | 8936 | 31494 |
| ATVLFRGGPRGLL | 7517 | 8908 |
| TVLFRGGPRGLLA | 7199 | 12668 |
| VLFRGGPRGLLAV | 236 | 628 |
| LFRGGPRGLLAVA | 24754 | 31034 |
| FRGGPRGLLAVAQ | 27133 | 32927 |
| RGGPRGLLAVAQD | 33051 | 37463 |
| GGPRGLLAVAQDL | 7960 | 30542 |
| GPRGLLAVAQDLT | 13423 | 30972 |
| PRGLLAVAQDLTE | 17551 | 31764 |
| RGLLAVAQDLTED | 16889 | 33306 |
| GLLAVAQDLTEDE | 17723 | 35216 |
| LLAVAQDLTEDEE | 34948 | 36072 |
| FPATVLFRGGPRGL | 9143 | 27652 |
| PATVLFRGGPRGLL | 7833 | 9280 |
| ATVLFRGGPRGLLA | 7941 | 14000 |
| TVLFRGGPRGLLAV | 351 | 958 |
| VLFRGGPRGLLAVA | 649 | 1403 |
| LFRGGPRGLLAVAQ | 27460 | 32988 |
| FRGGPRGLLAVAQD | 28291 | 34050 |
| RGGPRGLLAVAQDL | 7397 | 29261 |
| GGPRGLLAVAQDLT | 13582 | 31110 |
| GPRGLLAVAQDLTE | 17371 | 31898 |
| PRGLLAVAQDLTED | 19135 | 33912 |
| RGLLAVAQDLTEDE | 17765 | 34570 |
| GLLAVAQDLTEDEE | 18231 | 35941 |
| LLAVAQDLTEDEET | 24639 | 25997 |

## Transparent Methods

### Cell culture

HAP1 (HLA-A*02:01, HLA-B*40:01, HLA-C*03:04), EBV-LCLs, HEK293T, U266, U2G10196, COV413b, T2, UM9, and TMD8 cells were cultured in Iscove’s Modified Dulbecco’s Medium (IMDM, Gibco and Lonza) supplemented with 10% heat-inactivated fetal calf/bovine serum (Gibco, Thermo Fisher) and antibiotics (PenStrep; Invitrogen; IMDM^++^) at 37°C and 5% CO_2_. Primary HUVECs (kindly provided by Dr. Jaap van Buul), pooled from three different donors (Lonza), were cultured in Endothelial Cell Growth medium 2 (EGM2, Lonza) complemented with EGM2 Supplement mix (Promocel), 10% fetal calf serum and antibiotics. The HLA-A*02:01 restricted T cell clones were previously described and recognize a peptide derived from the endogenously expressed human proteins USP11, VPS13B or SSR1 (Amir et al., 2011; Van Bergen et al., 2010). The clones were expanded using irradiated feeder cells (mixed PBMCs and EBV-LCLs exposed to 30 Gy and 50 Gy, respectively) in IMDM with 5% fetal calf serum and 5% human serum (Sanquin), supplemented with 120 U/mL IL-2 (Miltenyi and Chiron), 0.8 μg/mL PHA (HA16 Remel™, ThermoFisher) and antibiotics as described (Pont et al., 2019; Spaapen et al., 2007).

### PBMC isolation and activation

Blood was withdrawn from 3 HLA-A2 positive healthy volunteers (Sanquin), PBMCs were isolated by Ficoll gradient separation. Monocytes were isolated through plastic-adherence by incubating fresh PBMCs in IMDM^++^ for 1 h at 37°C and 5% CO_2_ on culture plastic. B cells were isolated using anti-CD19 Dynabeads and DETACHaBEAD (Invitrogen) according to manufacturer’s protocol. B cell were activated by incubating them with 0.1 μM CpG (Invivogen) and 2.5 μg/mL Fab mix (anti-IGA, IgG and IgM, Jackson ImmunoResearch) in IMDM^++^ for 3 days at 37°C and 5% CO_2_.

### Sequencing

DNA was isolated from full PBMCs and HLA-A2 positively sorted HUVECs using the NucleoSpin Tissue kit (Machery-Nagel) according to manufacturer’s protocol. DNA was amplified using Taq DNA-polymerase (VWR) with forward primer TTCTTCGGTGCATTGTAATTG and reverse primer GGGTCAATGAAACCTTTTTCC and subsequently Sanger sequenced using the forward primer or a sequencing primer GGTGAACTGGGAGATTCTGTT (HUVECs) and BigDye v1.1 (Applied Biosystems).

### Genome-editing

Genome-editing and verification was performed as previously described for TAP1 (de Waard et al., 2020). In short, cells were transduced with pL.CRISPR.efs.GFP (Addgene) containing a gRNA targeting TAP1 (ACTGCTACTTCTCGCCGACT). Transduced cells were enriched by FACS sort on GFP positivity and clones were subsequently generated by limiting dilution. The targeted region was sequenced using primer (GCTCCCCATGAGATCAGCTC) after PCR on genomic DNA of single cell-derived clonal cell lines using forward primer (CAGCCTGTTCCTGGGACTTT) and reverse primer (ACTGACAACGAAGGCGGTAG).

### T cell assays

50,000 or 40,000 (only for activated B cells) target cells were cocultured with T cells in a 1:1 ratio in IMDM^++^ for 18 hours as previously described (Spaapen et al., 2008). IFN-γ and GM-CSF release was measured by standard sandwich ELISA (Sanquin and BioLegend, respectively) according to the manufacturer’s protocol. For IFN-γ stimulation, target cells were seeded two (HAP1 cells, HUVECs) or three (PBMCs) days prior to the experiment in a culture flask in IMDM^++^ containing 20 U/mL recombinant human IFN-γ (Peprotech).

### Flow Cytometry

Trypsinized cells were incubated with antibodies W6/32-PerCP-eFluor710 (pan-HLA; eBioscience) or BB7.2-APC (HLA-A2; eBioscience) in PBS for 30 min at 4°C. Stained cells were fixed in PBS containing 1% formaldehyde (Merck) and DAPI (1 μM, Sigma-Aldrich) for direct analysis by flow cytometry. Samples were analyzed on BD flow cytometers (LSRFortessa or LSR-II). FACS data was analyzed using FlowJo (Tree Star, Inc). Gating strategy is shown in Figure S1. The pooled primary HUVECs were incubated with BB7.2-APC antibody for 30 min at 4°C before they were sorted into HLA-A2 positive and negative cell populations in the presence of DAPI (1 μM) on a BD FACSAria II.

### Sample collection and HLA-I peptide elution

Peptide elution was performed as outlined previously (Hassan et al., 2013). Prior to cell pellet generation, UM9, and TMD8 were retrovirally transduced with HLA alleles to express HLA-A*02:01 (UM9-A2, TMD8-A2) and magnetic-activated cell sorting (MACS) enriched for nerve growth factor receptor (NGFR) marker gene expression using anti-R-phycoerythrin (PE) microbeads (Miltenyi Biotec, Bergisch Gladbach, Germany). Cell pellets were made of 5 primary acute myeloid leukemias (AML), 4 primary B cell malignancies (ALL, CLL, HCL), 2 primary ovarian carcinoma (1x ascites) and 2 multiple myeloma cell lines (U266 and UM9-A2), 1 DLBCL cell line (TMD8-A2) and the cell lines T2 and HAP1(de Waard et al., 2020). Primary ovarian carcinoma tumor material was first sliced into small pieces. Dead, clotted or non-tumor material was removed, small tumor tissue was put into a C-tube (Miltenyi Biotec) with ice cold buffer without detergent and complete protease inhibitor (Sigma-Aldrich) to prevent protein degradation and dissociated using a gentleMACS (Miltenyi Biotec) procedure until an almost homogenous cell solution. Cells were lysed in 50 mM Tris-HCl, 150 mM NaCl, 5 mM EDTA, and 0.5% Zwittergent 3-12 (pH 8.0) supplemented with complete protease inhibitor and Benzonase (Merck; 125 IU/mL to remove DNA/RNA complexes). After 2 hours of tumbling in lysis buffer at 4°C, cell lysates were centrifuged for 10 min at 1,000 g at 4°C. The supernatant was subsequently centrifuged for 35 min at 13,000 g at 4°C, precleared with Protein A Sepharose CL-4B beads (GE Healthcare Life Sciences), and subjected to an immunoaffinity column with dimethyl pimelimidate-immobilized W6/32 antibody (2-3 mg/ml resin) on Protein A Sepharose CL-4B beads. After flow through of the cell lysate, the column was washed with 5 to 10 column volumes of lysis buffer and 10 mM Tris-HCl (pH 8.0) buffers with 1 M, 120 mM, and no NaCl, bound HLA class I–peptide complexes were eluted from the column and dissociated with 3 to 4 column volumes of 10% acetic acid. Peptides were separated from HLA class I molecules via passage through a 10 kDa membrane (Macrosep Advance Centrifugal Devices with Supor Membrane, Pall Corp.) and freeze-dried.

### Fractionation and mass spectrometry (MS) of HLA-class I peptides

Eluted peptide pools were either fractionated by strong cation exchange chromatography (SCX) or by high pH reversed phase fractionation (High pH-RP). SCX fractionation was performed with a homemade 15 cm SCX column (320 μm inner diameter; polysulfoethyl A, 3 μm, Poly LC) run at 4 μL/min. Gradients were run for 10 min at 100% solvent A (100/0.1 water/trifluoroacetic acid v/v), after which a linear gradient started to reach 100% solvent B (65/35/0.1 250 mM KCl/acetonitrile (ACN)/trifluoroacetic acid v/v/v) over 15 min, followed by 100% solvent C (65/35/0.1 500 mM KCl/ACN/trifluoroacetic acid v/v/v) over the next 15 min. The gradient remained at 100% solvent C for 5 min and then switched again to 100% solvent A. Twenty 4 μL fractions were collected in vials prefilled with 20 μL 95/3/0.1 water/ACN/formic acid v/v/v and freeze-dried. High pH-RP fractionation was performed on an Oasis HLB 1cc SPE column (Waters). The column was first equilibrated with 1 mL 10/90 water/ACN v/v, washed 2 times by passing 1 mL of 10 mM ammonium bicarbonate (ambic) buffer pH=8.4 through the column. The freeze-dried peptide mixture was diluted in 10 mM ambic pH 8.4 and gently pushed through the column and washed again with 1 mL of 10 mM ambic pH 8.4. Fractions were taken by steadily increasing the amount of ACN into the elution buffer in the range of 5% to 50% ACN and freeze-dried.

Freeze-dried peptide fractions were dissolved in 95/3/0.1 water/ACN/formic acid v/v/v and analyzed as described previously (van der Lee et al., 2019). The peptides were analyzed by data-dependent MS/MS on a Q Exactive mass spectrometer equipped with an easy-nLC 1000 (Thermo Fisher Scientific) oran LTQ FT Ultra equipped with a nanoflow liquid chromatography 1100 HPLC system (Agilent Technologies). Peptides were trapped at 6–10 μL/min on a 1.5 cm column (100 μm internal diameter; ReproSil-Pur C18-AQ, 3 μm, Dr. Maisch HPLC GmbH) followed by elution to a 20 cm column (50 μm internal diameter; ReproSil-Pur C18-AQ, 3 μm) at 150 nL/min. The column was developed with a 0-40% ACN gradient in 0.1% formic acid for 120 min. The eluent was sprayed into the mass spectrometer by drawing the end of the column to a tip (internal diameter of ~5μm). Full-scan MS spectra were acquired in the FT-ICR with a target value of 3,000,000 at a resolution of 25,000. The 2 most intense ions were isolated for accurate FT-ICR mass measurements by a selected ion monitoring scan with a target accumulation value of 50,000 at a resolution of 50,000followed by fragmentation in the linear ion trap using collision-induced dissociation at a target value of 10,000. The Q Exactive mass spectrometer was used in top10 mode with an AGC target value of 3,000,000/maximum fill time of 20 ms (full scan) at a resolution of 70,000 and an AGC target value of 100,000/maximum fill time of 60 ms for MS/MS at a resolution of 17,500 and an intensity threshold of 17,000. Apex trigger was 1-10 sec with 2-6 allowed charges. The Orbitrap Fusion LUMOS mass spectrometer was used in data-dependent MS/MS (top-N mode), with recording of the MS2 spectrum in the orbitrap with a collision energy at 32 V. For master scan (MS1), the AGD target was 400,000/maximum fill time of 50 ms at a resolution of 60,000 and scan range 300-1,400. Charge states 1–4 were included and the dynamic exclusion was set after n=1 with a duration of 20 sec. For MS2, precursors were isolated using the quadrupole with an isolation width of 1.2 Da. HCD collision energy was at 32 V, first mass at 110 Da. The MS2 had an AGC target of 50,000/maximum fill time of 100 ms with a resolution of 30,000. To identify peptides and proteins, Proteome Discoverer version 2.1 (Thermo Fisher Scientific) was used with the mascot node for identification in mascot version 2.2.04 with the UniProt Homo Sapiens database (UP000005640; Jan 2015; 67,911 entries). Cysteine modifications were set as variably for Methionine oxidation and cysteinylation. Precursor tolerance of 10 ppm and MS/ MS fragment tolerance of 20 mmu was used for peptide assignment for the Q Exactive data and 2 ppm and 0.5 Da for LTQ FT Ultra data.

### Synthetic peptide stimulation

Synthetic peptides were titrated on 20,000 T2 cells prior to incubation with 5,000 T cells for 18 hours before cytokine ELISA similar as previously described (Bijen et al., 2018).

### Expression analysis

Data on SSR1, CD3E, CD14 and CD19 RNA expression in blood cells and of SSR1, CD45 and PECAM1 in HUVECs was retrieved from the Human Protein Atlas (https://www.proteinatlas.org)(Uhlen et al., 2015) using the RNA HPA blood cell gene and the RNA HPA cell line gene datasets. Data for B and T cells was pooled from the different naïve and memory subsets. Microarray data of IFN-γ stimulated human cells were selected based on equal *in vitro* treatment with 200 U/mL IFN-γ for 24 hours containing expression data on SSR1 (downloaded from Interferome.org v2.01, datasets 64, 144, 311). PPIA was selected as house-hold control as it is unaffected by interferon signaling and was present in all datasets (Riemer et al., 2012). VPS13B and USP11 expression was only present in datasets 64 and 144, respectively.

### Statistical Analyses

All error bars correspond to the standard deviation of the mean. Statistical evaluations were done by a student’s t-test for comparison of two independent groups (T cell assay on HAP1 wild type and TAP1 KO cells), a repeated measures one-way ANOVA followed by Dunnett’s multiple comparisons test for comparison of change in gene expression derived from multiple datasets (IFN-γ regulation of mRNA), an one-way ANOVA followed by Dunnett’s multiple comparisons test (IFN-γ regulation of protein) or Sidak’s multiple comparisons test (T cell assay on IFN-γ stimulated HAP1 cells) for multigroup analyses with Prism software (http://www.graphpad.com). Statistical evaluation of primary cell cocultures was unnecessary due to the qualitative goal of the experiments.

